# A Phased *Canis lupus familiaris* Labrador Retriever Reference Genome Utilizing High Molecular Weight DNA Extraction Methods and High Resolution Sequencing Technologies

**DOI:** 10.1101/2020.08.26.269076

**Authors:** Robert A. Player, Ellen R. Forsyth, Kathleen J. Verratti, David W. Mohr, Alan F. Scott, Christopher E. Bradburne

## Abstract

Reference genome fidelity is critically important for genome wide association studies (GWAS), yet many are incomplete or too dissimilar from the study population. A typical whole genome sequencing approach implies short-read technologies resulting in fragmented assemblies with regions of ambiguity low complexity. Further information is lost by economic necessity when genotyping populations, as lower resolution technologies such as genotyping arrays are commonly utilized. Here we present a phased reference genome for *Canis lupus familiaris* utilizing high molecular weight sequencing technologies. We tested wet lab and bioinformatic approaches to demonstrate a minimum workflow to generate the 2.4 gigabase genome for a Labrador Retriever. The resulting *de novo* assembly required eight Oxford Nanopore R9.4 flowcells (~23X depth) and running a 10X Genomics library on the equivalent of one lane of an Illumina NovaSeq S1 flowcell (~88X depth), bringing the cost of generating a nearly complete reference genome to less than $10K. Mapping of publicly available short-read data from ten Labrador Retrievers against this breed-specific reference resulted in an average of approximately 1% more aligned reads compared to mapping against the current gold standard reference (CanFam3.1, p<0.001), indicating a more complete breed-specific reference. An average 15% reduction of variant calls was observed from the same mapped data, which increases the chance of identifying low effect size variants in a GWAS. We believe that by incorporating the cost to produce a full genome assembly into any large-scale canine genotyping study, an investigator can make an informed cost/benefit analysis regarding genotyping technology.

## INTRODUCTION

The revolution in genomic sequencing technologies is creating a wealth of information about diverse taxa. Typically, an organism is sequenced as a high quality reference, and then the variability in genomic content within individuals is surveyed using cheaper, more economically viable technologies (Green and Guyer 2011). Over time, the costs of genomic characterization are reduced as technological performance increases. This means that periodically, new references need to be established that can be used for read mapping and scaled genotyping approaches, such as the design of new Single Nucleotide Polymorphism (SNP) arrays used to genotype large numbers of individuals. An example is the human genome, which was established in draft form in 2001 at a cost of $3.2B US (Venter et al. 2001). Following completion, haplotyping of populations continued at a large scale using high-throughput SNP chips, which initially started with a few hundred thousand SNPs but within 10 years contained millions. Likewise the human reference has been continually updated, starting in 2001, with a draft sequence covering more than 90% of the genome, had a 1:1000 base pair (bp) error rate, and contained 150,000 gaps. Within two years the same genome had reached 99% coverage, 1:10,000 bp error rate, and only 400 gaps (“Human Genome Project FAQ” n.d.). According to the National Human Genome Research Institute (NHGRI) tracking site, the cost has stabilized at around $1K per full human genome since 2015.

However, the human genomes considered for this estimation do not come close to full completion, having a 1:100 bp error rate along with widely varying percent coverage (“DNA Sequencing Costs: Data” n.d.). The $1K estimate also assumes the utilization of whole genome sequencing (WGS) short read technologies. For Genome-Wide Association Studies (GWAS), lack of genetic information due to incomplete genomes can lead to false negatives from an inability to see real variants, or false positives from false variant calls against a reference. In fact, the early reliance on SNPs to type the variation in humans has likely contributed to the ‘missing heritability’ problem of human genomic medicine (Manolio et al. 2009; Young 2019).

Canids share a similar story. The current reference sequence for canids is a boxer: CanFam3.1, submitted to NCBI in November of 2011 (Kim et al. 1998; Lindblad-Toh et al. 2005). It was sequenced with Illumina short read technologies and has been continuously updated ever since (the latest update as of this article was in June of 2019) (“Canis Lupus Familiaris - Ensembl Genome Browser 100” n.d.). Various SNP genotyping chips, whose costs are dependent on scale but average $100-$500 per animal, have been developed, but much of the detectable genetic variation depends on an incomplete and constantly changing reference. Long read technologies have the potential to change this paradigm and lead the community to generate single reference genomes for individual projects. The longer read lengths of approximately 2 to 30 kb (kilobase) remove many of the bioinformatic challenges inherent in short read sequencing and allow previously unheard of resolution to observe structural variants and the organization of long stretches of low-complexity DNA A genome assayed with this ‘high-resolution genomic’ approach using longer reads could provide structural variants together with SNPs. Further, application of high-resolution genomics across a population for a GWAS could illuminate any ‘missing heritability’ for a population, such as structural variants that are unresolvable with SNP or WGS short read platforms. Canids provide an excellent test case for this approach.

*Canis lupus familiaris* has been under selection by human breeding for thousands of years, which has created extremely variable morphologies within a single species (Plassais et al. 2019). Therefore, unlike human genomes that have many common variants of low effect size, dogs have many common variants of large effect size. Any study that lacks genomic context of a breed by not having a high-quality reference genome specific to that breed runs the risk of missing important SNPs and structural variants that may be associated with interesting phenotypes. We set out to establish the best workflows to provide the highest quality genome at the lowest cost, taking advantage of Oxford Nanopore Technologies (ONT), 10X Genomics, and Illumina sequencing technologies. The resulting genome is of a yellow Labrador Retriever, named ‘Yella’, and we estimate that similar workflows could be used to easily generate high - quality reference genomes for researchers or breeders establishing studies requiring high-resolution variation. Further, we assert that any large-scale study on genetic variation for a population should begin with the establishment of a local high-quality reference genome for that population.

## RESULTS

When setting out to produce a high-quality, phased reference genome, careful consideration should be given to wet lab processes that do the following: 1) provide optimal preservation for downstream extraction, 2) generate high quantity and quality of high molecular weight (HMW) DNA, and 3) are robust and reproducible (i.e., they provide the least amount of variability between different individual blood samples). Figure 1 shows the wet lab process flow and components that were evaluated in this study, and used to generate HMW canine DNA for sequencing and *de novo* genome assembly.

**Figure 1.**
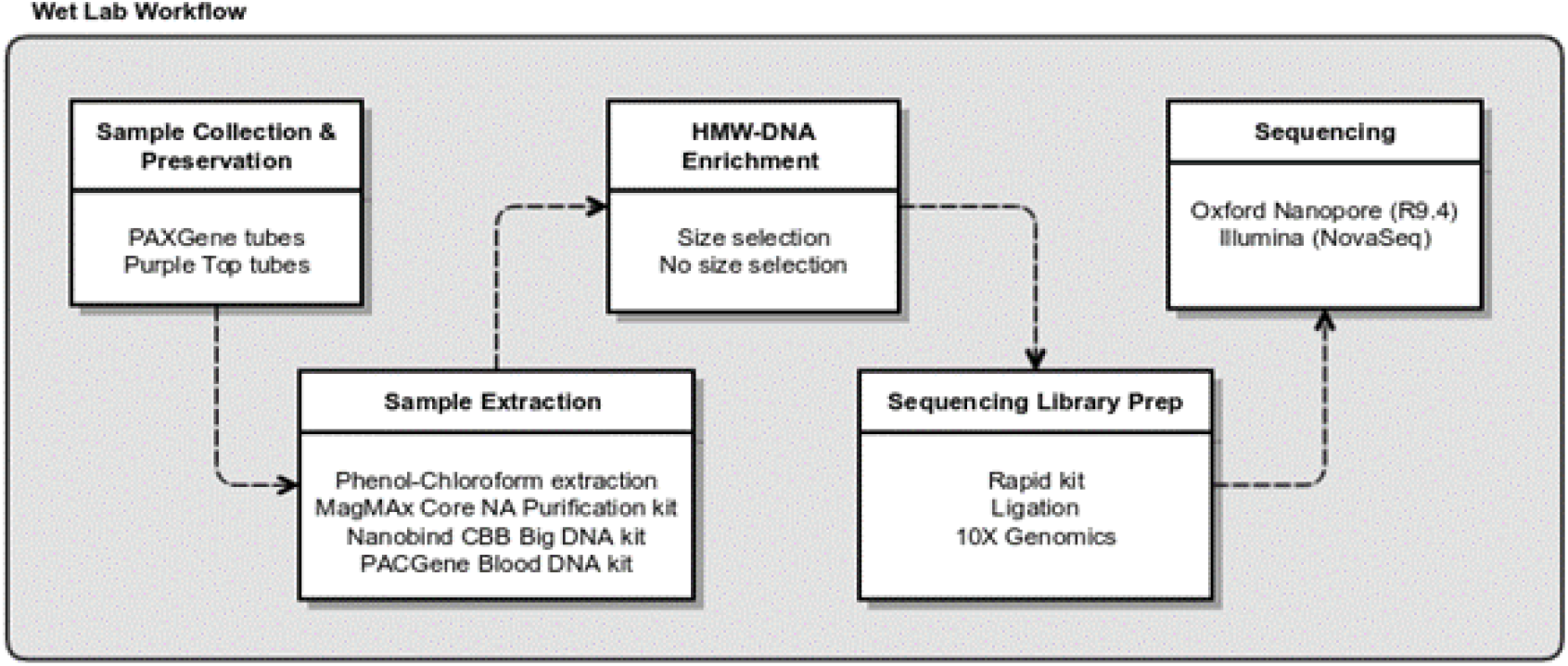
Diagram of wet lab workflow for testing sample collection, extraction, and sequencing library preparation methods used in this study.

### Preservation, extraction, and acquisition of HMW-DNA

Canine blood samples were collected and delivered in either the PAXgene DNA proprietary storage media or a purple top Vacutainer tube with EDTA (ethylenediaminetetraacetic acid). These two preservative types were evaluated in conjunction with four DNA extraction and isolation methods: 1) a standard phenol chloroform extraction (PCE) method, 2) the Magmax Core NA Purification, 3) the Nanobind CBB Big DNA kit, and 4) the PAXgene Blood DNA kit. Blood samples from Yella stored in the purple top tubes and extracted with the Nanobind kit yielded the best purity (highest 260/280 ratio) and highest concentrations (Table 1, additional information in Table S1). Compared to PCE from the same storage method, this is equivalent to a 92-fold increase in extraction efficiency. In terms of total recovered NA, the PAXgene extraction from the purple top tube performed best, yielding over 10 ug DNA Most importantly, significant fractions of HMW-DNA using the PAXgene extraction kit were not detected (Figure S1). Direct comparison of extraction kits showed that the Nanobind kit provided the most consistent DNA yield and quality among the four kits tested using blood stored in EDTA from four different canines (Table 2 and Figure 2).

**Figure 2.**
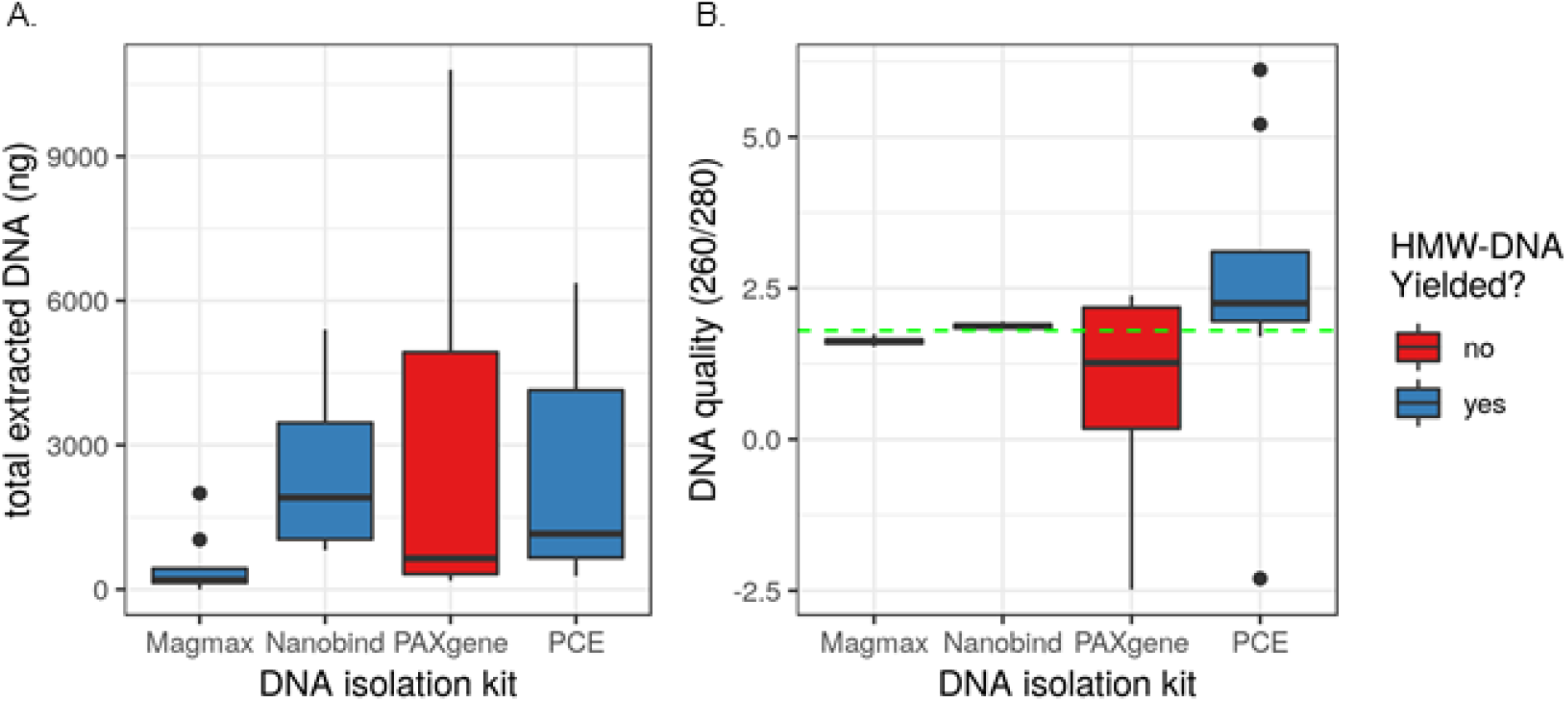
Total extracted DNA and DNA quality from four tested isolation kits. A) Total extracted DNA. B) DNA quality; green line indicates the ideal 260/280 ratio for DNA purity at 1.80. Extractions from the Nanobind kit had the most consistently high yield and quality.

**Table 1.** Effect of blood sample preservation agent on DNA yield. Blood for one canine (Yella) was drawn directly into two tubes containing either a proprietary preservation agent, or EDTA. Three kits were tested against a phenol-chloroform extraction (PCE) standard method. Input and output volumes for each kit are shown, along with actual recovered total DNA mass. NA stands for nucleic acid. EDTA stands for ethylenediaminetetraacetic acid.

| Storage Agent | NA Isolation Method | Input Volume (uL) | Output Volume (uL) | NA Conc (ng/uL) | Recovered NA Total (ng) | NA Quality (260/280) | HMW DNA Yielded? |
| --- | --- | --- | --- | --- | --- | --- | --- |
| proprietary (PAXgene) | PCE | 1700 | 1000 | 6.37 | 6370 | 2.20 | yes |
| proprietary (PAXgene) | Magmax Core NA Purification | 200 | 90 | 2.03 | 183 | 1.66 | yes |
| proprietary (PAXgene) | Nanobind CBB Big DNA kit | 200 | 100 | 11.10 | 1110 | 1.87 | yes |
| proprietary (PAXgene) | PAXgene Blood DNA kit | 1700 | 1000 | 6.40 | 6400 | 2.38 | no |
| EDTA (purple top) | PCE | 1700 | 1000 | 0.38 | 380 | 5.21 | yes |
| EDTA (purple top) | Magmax Core NA Purification | 200 | 90 | 2.63 | 237 | 1.62 | yes |
| EDTA (purple top) | Nanobind CBB Big DNA kit | 200 | 100 | 35.30 | 3530 | 1.84 | yes |
| EDTA (purple top) | PAXgene Blood DNA kit | 1700 | 1000 | 10.80 | 10800 | 1.98 | no |

**Table 2.** Variability of NA (nucleic acid) isolation method across four canine blood samples preserved in ‘purple top’ tubes with EDTA (ethylenediaminetetraacetic acid). DNA from purple top tubes was extraced using either phenol-chloroform extraction (PCE), or three commercial kits (Magmax, Nanobind, and PAXgene). Bold values represent the best performance in a particular category.

| NA Isolation Method | Total NA (ug) Mean | Total NA Std. Dev. | NA Quality (260/280) Mean | NA Quality (260/280) Std. Dev. | High-MW DNA? |
| --- | --- | --- | --- | --- | --- |
| PCE | 1.28 | 1.63 | 1.75 | 3.10 | yes |
| Magmax Core NA Purification | 0.81 | **1.02** | 1.57 | 0.05 | yes |
| Nanobind CBB Big DNA kit | 2.92 | 1.99 | **1.85** | **0.03** | yes |
| PAXgene Blood DNA kit | **4.08** | 4.85 | 1.73 | 0.79 | no |

### DNA Size-selection and Oxford Nanopore Sequencing

Estimated average genome depth, based on the 2.32 Gb (gigabase) CanFam3.1 genome, for combined read data from all eight ONT R9.4.1 flow cells was 22.65x (Table 3). Additional read statistics for the combined read data are shown in Figure S2. The read N50 varied per flow cell dataset from 11,868 to 35,584 bp (Table 4). Interestingly, size selection with the Circulomics Short Read Eliminator kit prior to library preparation did not always result in a higher read N50, and in fact the read N50 was actually reduced when the kit was used prior to library preparation with the ligation kit (SQK-LSK109). Instead, read N50 appears more influenced by library kit type, with the ligation kit having approximately 2x higher median read N50 than the rapid kit (SQK-RAD004) (median read N50 of 24,750 and 12,094 bp, respectively).

**Table 3.** Breakdown of ONT sequencing runs, flow cells, library kit type, and estimated depth shown in Figure 1. Flow cell number from Table 2.

| Run # | Flowcell # | ONT kit | Total flow cells | Est. depth |
| --- | --- | --- | --- | --- |
| 1 | 1,2 | SQK-LSK109 | 2 | 6.66 |
| 2 | 5,6 | SQK-LSK109 | 2 | 3.96 |
| 1+2 | 1,2,5,6 | SQK-LSK109 | 4 | 10.02 |
| 1 | 3,4 | SQK-RAD004 | 2 | 5.99 |
| 2 | 7,8 | SQK-RAD004 | 2 | 6.65 |
| 1+2 | 3,4,7,8 | SQK-RAD004 | 4 | 12.64 |
| 1 | 1,2,3,4 | RAD+LSK | 4 | 12.05 |
| 2 | 5,6,7,8 | RAD+LSK | 4 | 10.6 |
| 1+2 | 1,2,3,4,5,6,7,8 | RAD+LSK | 8 | 22.65 |

**Table 4.**
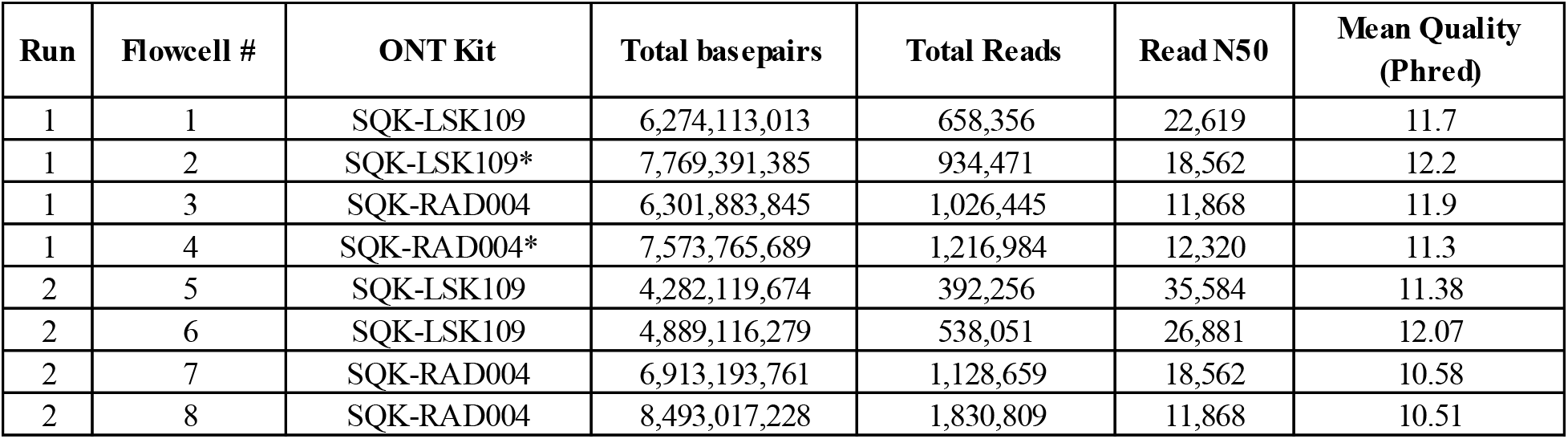
Oxford Nanopore GridION sequencing run summaries using R9.4.1 flowcells. SQK-LSK109 is the ligation based library preparation kit. SQK-RAD004 is the transposon based rapid library preparation kit. * Size selection on extracted DNA, prior to library preparation using the Circulomics short read eliminator kit.

| Run | Flowcell # | ONT Kit | Total basepairs | Total Reads | Read N50 | Mean Quality (Phred) |
| --- | --- | --- | --- | --- | --- | --- |
| 1 | 1 | SQK-LSK109 | 6,274,113,013 | 658,356 | 22,619 | 11.7 |
| 1 | 2 | SQK-LSK109* | 7,769,391,385 | 934,471 | 18,562 | 12.2 |
| 1 | 3 | SQK-RAD004 | 6,301,883,845 | 1,026,445 | 11,868 | 11.9 |
| 1 | 4 | SQK-RAD004* | 7,573,765,689 | 1,216,984 | 12,320 | 11.3 |
| 2 | 5 | SQK-LSK109 | 4,282,119,674 | 392,256 | 35,584 | 11.38 |
| 2 | 6 | SQK-LSK109 | 4,889,116,279 | 538,051 | 26,881 | 12.07 |
| 2 | 7 | SQK-RAD004 | 6,913,193,761 | 1,128,659 | 18,562 | 10.58 |
| 2 | 8 | SQK-RAD004 | 8,493,017,228 | 1,830,809 | 11,868 | 10.51 |

### Illumina Sequencing of 10X Genomics Library and SuperNova Scaffolding

Estimated average genome depth for trimmed reads data from four lanes of Illumina NovaSeq was 87.80x (Table 5). SuperNova scaffolding was performed, which utilizes the 10X GEM barcoding preparation for more accurate localization of short reads into contigs, under the assumption that reads sharing the same barcode are derived from the same small number of HMW DNA fragments contained in each GEM. The resulting scaffold contained 10,391 contigs, with a contig N50 and L50 of 94 kb and 22 contigs, respectively. The phase block size was greater than 5 Mb (megabase), and the scaffold N50 was 39 Mb. The assembly size of scaffolds greater than or equal to 10 kb was 2.33 Gb, which is in agreement with other canine breed assemblies such as the Boxer (CanFam3.1 assembly at 2.31 Gb) and German Shepherd (GCA_008641245.1 assembly at 2.36 Gb).

**Table 5.** Illumina 10X library, 300 cycle sequencing run summaries. Insert size ~400 bp, these libraries were not prepared with the intention of joining (hence the 100bp gap between pairs). Quality and adapter trimming was performed with cutadapt (including clipping the first 22 bases from R1).

| Run | Lane | Paired Read | RAW |  | TRIMMED |  |
| --- | --- | --- | --- | --- | --- | --- |
|  |  |  | Total bps | Total reads | Total bps | Total reads |
| 3 | 1 | 1 | 2.36E+10 | 156,607,429 | 2.00E+10 | 155,880,038 |
| 3 | 1 | 2 | 2.36E+10 | 156,607,429 | 2.35E+10 | 155,880,038 |
| 3 | 2 | 1 | 2.29E+10 | 151,709,875 | 1.94E+10 | 151,035,675 |
| 3 | 2 | 2 | 2.29E+10 | 151,709,875 | 2.27E+10 | 151,035,675 |
| 4 | 1 | 1 | 3.16E+10 | 209,187,620 | 2.68E+10 | 208,419,758 |
| 4 | 1 | 2 | 3.16E+10 | 209,187,620 | 3.14E+10 | 208,419,758 |
| 4 | 2 | 1 | 3.24E+10 | 214,451,964 | 2.75E+10 | 213,618,769 |
| 4 | 2 | 2 | 3.24E+10 | 214,451,964 | 3.22E+10 | 213,618,769 |
| **Totals** |  |  | 2.21E+11 | 1,463,913,776 | 2.03E+11 | 1,457,908,480 |
| **Est. depth** |  |  | 95.38 |  | 87.80 |  |

### De Novo Assembly

The effect of estimated average read depth and library preparation kit (SQK-RAD004 or SQK-LSK109, i.e. rapid or ligation, respectively) on assembly contig count and total length was examined. The overriding factor for achieving the expected ~2.35 Gb assembly length is read depth, with the combination of reads from all eight flow cells achieving the expected length and about a magnitude reduction in total contigs compared to the CanFam3.1 assembly. ONT kit type had less of an effect on total length and contig count, with the ligation-only assemblies (at 10.02x depth) achieving a higher total length than the rapid-only assemblies (at 12.64x depth), even at ~2.5x lower estimated depth. However, the ligation kit assemblies appear more influenced by miniasm parameter selection compared to the rapid kit assemblies. A combination of kit types at a similar estimated depth (12.05x) seems to be the best of both worlds, with resulting assemblies having approximately the same number of contigs as the rapid-only assemblies (i. e. lower than ligation-only assemblies) at a comparable total length to the ligation-only assemblies.

Next, the effect of parameters available in the *de novo* assembler called miniasm on the ~23x estimated genome depth assemblies was assessed by examining the assembly cluster at the top left of Figure 3. Figure 4 shows 144 assemblies, which correspond to 144 unique parameter sets tested. It is important to note, however, that since the ‘m’ parameter had no effect on the assembly attributes of interest, there appears to be only 48 points in each plot. The following correlations and description of effect on assembly attributes is with respect to an increasing parameter value (see Fig 3 legend for description of parameters): m, not correlated, no effect; i, negative correlation, slightly less total bps but more contigs; s, negative correlation, significantly less total bps but more contigs; I, positive correlation, moderately more contigs and total bps; e, positive correlation, less contigs and less total bps.

**Figure 3.**
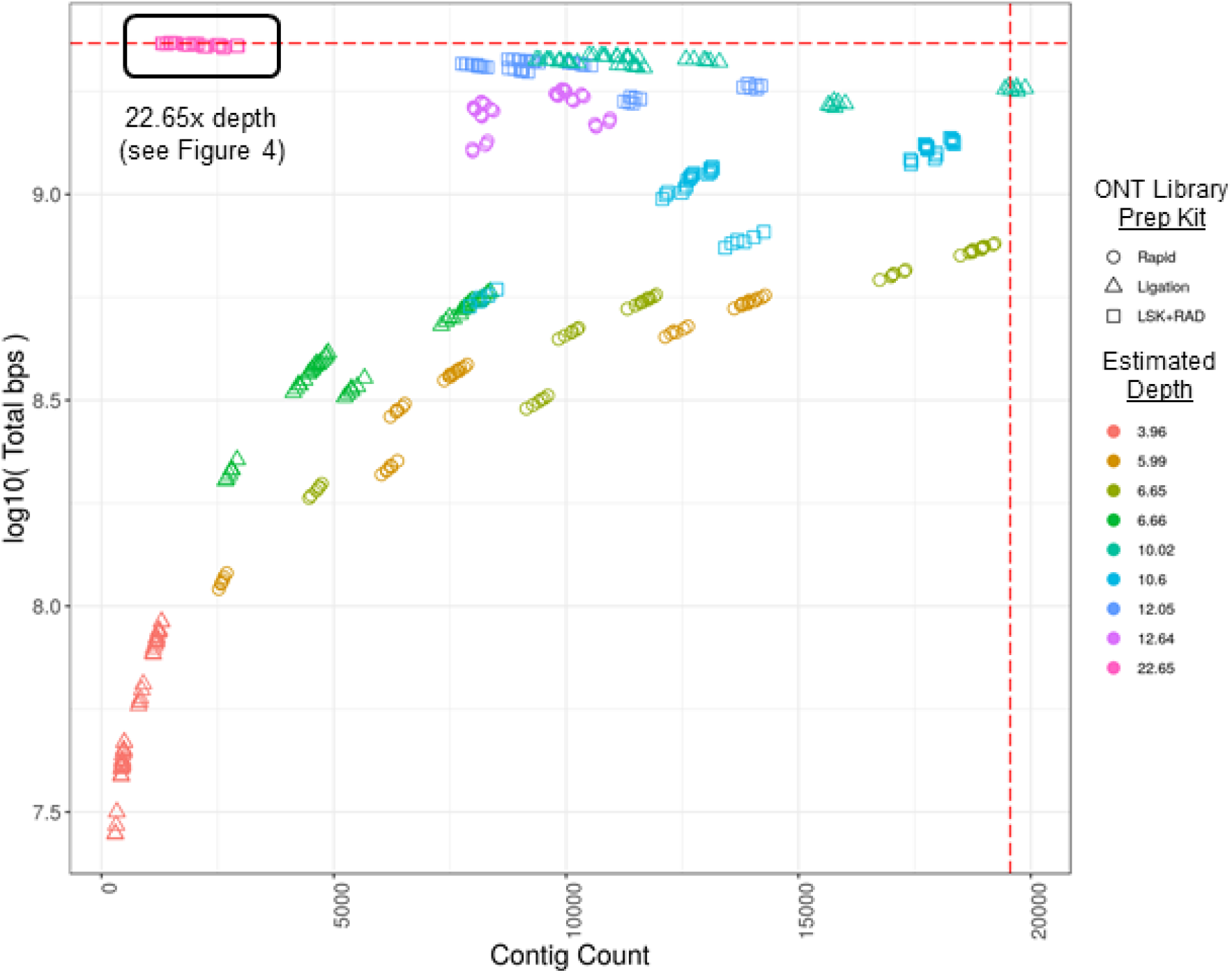
Genome assembly contig count versus total length of assembly. Each point represents a distinct assembly resulting from one of 144 unique miniasm parameter combinations. Sequence data from eight ONT flow cells are represented in the plot, four from each of the Ligation and Rapid library preparation kits (SQK-LSK109 and SQK-RAD004, respectively). See Table 3 for details linking ‘estimated depth’ to sequencing run and library kit. The estimated depth of 22.65 is a combination of reads from all eight flow cells (black boxed region in upper left, see Figure 4 for details regarding parameters). Estimated coverage is based on the total bps in the read set divided by the total length of CanFam3.1 assembly including Ns. Total bps of assembly approaches estimated total genome size as depth approaches 20x. Horizontal dashed red line - size of CanFam3.1 with N’s (2,327,604,993 bp); vertical dashed red line - contig count (19,555) of CanFam3.1 chromosomal scaffolds broken at every occurrence of N. The following ‘Estimated Depth(s)’ are from: the rapid kit only (5.99, 6.65, and 12.64); the ligation kit only (3.96, 6.66, 10.02); and a combination of the two (10.60, 12.05, and 22.65).

**Figure 4.**
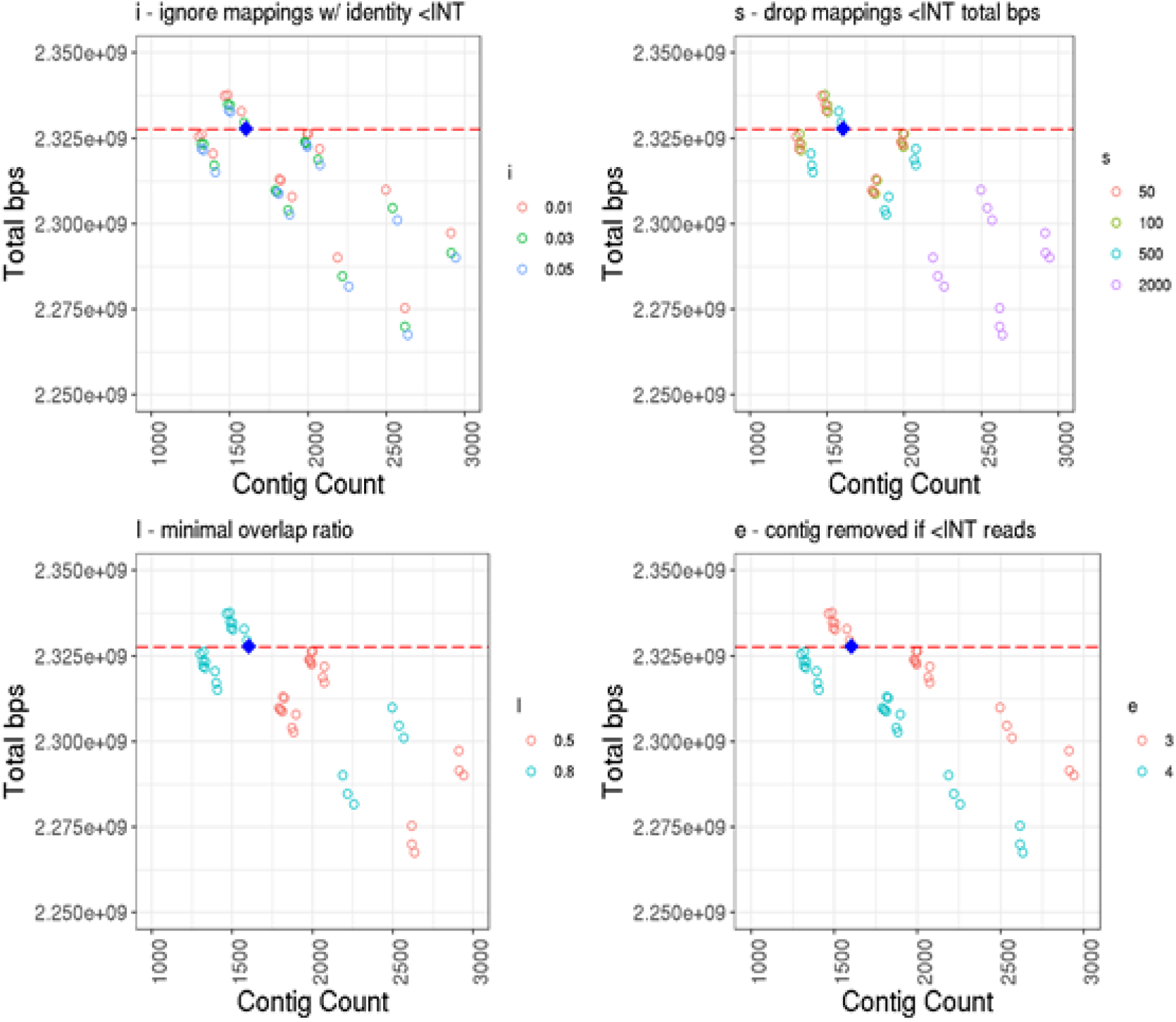
Genome assembly contig count versus total length of assembly for 22.65x estimated genome depth data. Contig count calculated from counting number of headers in resulting assembly FASTA files, and total length calculated from non-header character count. Zoomed in view of the top-left group of assemblies from Figure 3, colored by parameter value and broken down by miniasm parameter type: i, ignore mappings with identity less than INT (integer) identity; s, drop mapping less than INT total bps; I, minimap overlap ratio; and e, contig is removed if it is generated from less than INT reads. Note that miniasm parameter ‘m’ (for dropping read mappings with less than INT matching bps) is left out, as all points for the three values used (25, 50, and 100) are all overlapping (i.e. ‘m’ has no effect on contig count or total bps). Default parameters for miniasm are: m=100, i=0.05, s=1000, I=0.8, e=4. The blue diamond indicates the down-selected assembly (v0.0 in Table 4a) used for polishing and final scaffolding, miniasm parameters used: m=100, i=0.05, s=500, I=0.8, e=3. The red dashed line indicates the genome size (with N’s) of CanFam3.1.

The miniasm parameters used for the down-selected assembly that was subsequently polished and used for genome scaffolding (Table 6, v0.0) were ‘ -m 100 -i 0.05 -s 500 -I 0.8 -e 3’. These settings are only slightly less stringent than the default settings (-m 100 -i 0.05 -s 1000 -I 0.8 -e 4), with mappings less than 500 instead of 1000 total bases dropped (-s), and contigs generated from less than 3 instead of 4 reads removed (-e). The three parameters that remained at the default value are all more stringent compared to other parameter set values tested. The assembly was selected based on its relatively low contig count compared to that produced from other parameters sets, and a total assembly length approaching that of CanFam3.1.

**Table 6.** Assembly metrics of Yella dog genome through the scaffolding process, with related dog genome assembly metrics for comparison. BUSCO scores calculated using v3 with the mammalia_odb9 dataset (missing % equals 100 - [Complete+Fragmented]).

| Description | Total Contigs | Largest Contig | Total Length (Gb) | GC Content | N50 (Mb) | L50 | N per 100Kb | BUSCO Scores |  |
| --- | --- | --- | --- | --- | --- | --- | --- | --- | --- |
|  |  |  |  |  |  |  |  | Complete | Fragmented |
| CF, GCF_000002285.3 | 82 | 123,773,608 | 2.328 | 41.06% | 47.7 | 19 | 429 | 95.20% | 2.50% |
| GS, GCA_008641245.1 | 40 | 126,700,074 | 2.367 | 41.21% | 64.5 | 14 | 236 | 93.70% | 3.40% |
| CFGS, RaGOO of CF onto GS | 40 | 123,868,242 | 2.328 | 41.06% | 64.2 | 14 | 430 | 92.90% | 3.80% |
| JHMI 10X pseudohap | 10,391 | 96,528,903 | 2.417 | 41.25% | 39.2 | 22 | 1,901 | 92.70% | 4.40% |
| v0.0 | 1,601 | 20,780,228 | 2.299 | 41.11% | 5.5 | 130 | 0 | 0.20% | 1.10% |
| v0.1 | 1,600 | 21,039,211 | 2.326 | 40.98% | 5.6 | 130 | 0 | 32.00% | 21.80% |
| v0.2 | 1,600 | 21,018,819 | 2.324 | 41.17% | 5.6 | 130 | 0 | 94.80% | 2.70% |
| v0.3a | 1,412 | 21,088,418 | 2.394 | 41.30% | 5.4 | 134 | 270 | 95.20% | 2.60% |
| v0.3b | 1,413 | 21,084,388 | 2.394 | 41.30% | 5.4 | 134 | 270 | 95.20% | 2.50% |
| v0.4 | 40 | 131,668,473 | 2.435 | 41.30% | 64.9 | 14 | 1,972 | 92.40% | 4.20% |
| v1.0a | 40 | 138,659,542 | 2.394 | 41.30% | 64.3 | 14 | 276 | 95.00% | 2.50% |
| v1.0b | 40 | 138,666,786 | 2.493 | 41.30% | 64.3 | 14 | 276 | 95.10% | 2.30% |

Subsequent polishing of the v0.0 assembly using Racon resulted in large increases in ‘BUSCO complete’ percentages, starting at only 0.20% in v0.0 (unpolished assembly), 32.00% in v0.1 (3x ONT polishing), and 94.80% in v0.2 (2x Illumina polishing). After contig-level scaffolding of 10X contigs of each haplotype onto v0.2, then chromosome-level scaffolding of each v0.3 haplotype onto the v0.4 scaffold, BUSCO complete percentages were further increased to 95.00% and 95.10% for v1.0a and v1.0b, respectively (Table 6). These values are comparable to those achieved by CanFam3.1 at 95.20%. Compared to the 10X SuperNova pseudohap assembly the N per 100 kb metric was much improved through scaffolding onto the polished ONT scaffold (v0.2), from 1,901 down to only 275.90 and 275.77 in the final assembly haplotypes v1.0a and b, respectively. This suggests that the contiguous regions of the final assembly haplotypes are similar, the only differences being SNPs and small indels. Additionally, the CanFam3.1 reference contains 429 N per 100 kb, significantly more than the v1.0 assembly. Although the German Shepherd assembly (GCA_008641245.1) contains only 236 N per 100 kb, it only contains 93.7% complete BUSCOs. Overall, the total length of v1.0a and v1.0b are similar, at approximately 2.39 Gb, with the largest contig about 10% larger than that of either CanFam3.1 or the German Shepherd assembly.

### Mapping available public sequence data against reference genomes

In order to evaluate performance as a new reference genome, publicly available Illumina WGS reads from ten LRs were obtained from NCBI’s Sequence Read Archive (SRA). These are part of a 722 canid dataset, each sequenced with Illumina WGS and deposited on SRA in 2018 (accessions available in Table S2). It is one of the first datasets to be available for researchers to explore genomic variability among canid species beyond SNP-chip-level variation (Plassais et al. 2019). Ten Labrador Retriever data sets were mapped against three different canid breed reference genomes: Boxer (CF, CanFam3.1, GCF_000002285.3), German Shepard GS, GCA_008641245.1), and the Labrador Retriever genome presented here (YA, Yella_v1.0a, CP050567-CP050606). Figure 5 shows alignment rates and total high-quality variants called for each. In comparison to the Boxer and German Shepherd reference genomes, significantly more reads map to our Labrador Retriever reference, as expected (Figure 5A, paired Student’s t-test; CF vs YA p-value = 2.457e-06, GS vs YA p-value = 1.397e-03). One area in which a breed-specific reference would be expected to excel is when calling variants. Assuming that a genome specific to a breed has the most conserved structural and SNP variation, the number of called variants should decrease when reads from the same breed are mapped versus reads derived from a different breed. This can clearly be seen in Figure 5B, which shows the number of high-quality variants called (those with Q-score ≥ 30) from the ten Labradors mapped against each reference. Interestingly, the Boxer and Shepherd show similar performance when compared to total variants called in the Labrador, with the Labrador resolving an average of approximately15% of variants called against the non-Labrador breeds (Table S2).

**Figure 5.**
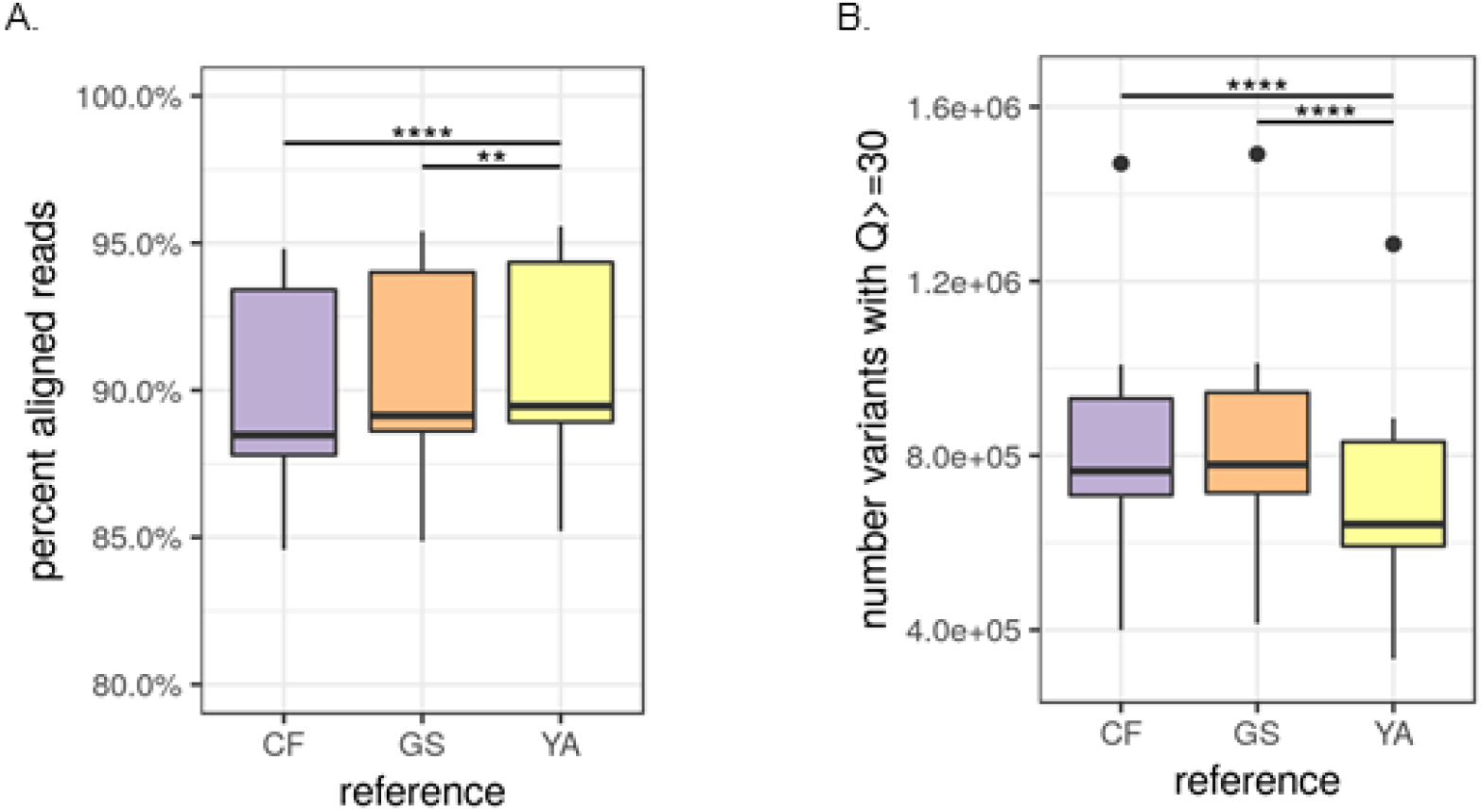
Alignment rates and total variants of ten Labrador Retriever Illumina sequence read data sets from SRA. Accessions and additional metrics can be found in Table S2. A) Reads alignment rates to CF (GCF_000002285.3, CamFam3.1, Boxer breed), GS (GCA_008641245.1, German Shepherd breed), and YA (Yella v1.0, Labrador Retriever breed) reference genomes (paired Student’s t-test; CF vs YA p-value = 2.457e-06, GS vs YA p-value = 1.397e-03). B) Total variants detected at Q-score ≥ 30 in references (paired Student’s t-test; CF vs YA p-value = 4.744e-06, GS vs YA p-value = 3.931e-06).

### Mitochondrial sequence and Y-chromosome

The mitochondrial (MT) genome was easily recoverable from Yella and comparable to the CanFam3.1 MT reference (Figure S3). It was annotated and visualized using GeSeq (Tillich et al. 2017). The Y-chromosome was much more recalcitrant. Yella is a male Labrador Retriever, and while reads from the Y-chromosome could be detected via alignment to an existing partial Y chromosome reference sequence, the Y-chromosome for Yella was not able to be resolved beyond an acceptable threshold for a published reference genome. This is similar to issues experienced across mammalian genomics, in which the short and highly repetitive nature of the Y-chromosome, along with its homology to the X-chromosome can make it difficult to detect and assemble (G. Li et al. 2013; Oetjens et al. 2018; Carvalho and Clark 2013; Rangavittal et al. 2019).

## DISCUSSION

Over the past two decades, much of the population-wide haplotyping of humans and dogs necessitated using SNPs derived from a single reference genome. In both cases, the starting references (a European American and a Boxer, respectively) would not be useful for ethnic stratification (for humans) or breed stratification (for canids). This can lead to an influx of false positives and false negatives when calling variants for a mixed population. In addition, the reliance on SNPs has failed to capture structural variation among populations, which has also not been well captured by array methodologies. One way to address both of these issues is the generation of a ‘stratified reference’ with cheaper technologies, such as short-read WGS, prior to initiating a GWAS. Here we provide the wet lab and bioinformatic methodology to generate a high-resolution mammalian reference genome for approximately $10K. Offsetting these costs would be the improved resolution of individuals mapped to the reference, and the elimination of a large proportion of variant call noise. We show that publicly-available canids generated with WGS can be re-mapped, allowing more comparative controls to be utilized for a GWAS without further expenditure. Investigators using this approach could affordably generate a high-quality GWAS using a high-resolution, stratified reference, and a population genotyped using WGS. In canids, this could allow for breed-specific elucidation of structural variants, and, more importantly, the determination of their frequencies within that breed. As frequencies of SNPs and structural variants are combined, this data could then be applied towards the ultimate genomic reference goal: the *Canis lupus familiaris* pan-genome.

## METHODS

### Sample collection

Blood samples were obtained from four canines, and collected in both PAXGene Blood DNA tubes (761115, PreAnalytix) and ‘purple top’ EDTA (ethylenediaminetetraacetic acid) Vacutainer tubes (367863, BD Biosciences). Blood samples were stored at 4C upon arrival and processed within 2 days. Samples were split between four different DNA extraction protocols (described below) to test extraction efficiency.

### DNA extraction and analysis of HMW-DNA

Four DNA extraction protocols were used to process blood samples: (1) the Dog Genome Project Protocol (“Online Research Resources Developed at NHGRI” n.d.) which employs a phenol-chloroform extraction (PCE), (2) the PAXgene Blood DNA kit (761133, PreAnalytix), (3) the MagMax Core NA kit (A32700, Applied BioScience), and (4) the Nanobind CBB Big DNA Kit (Beta Ultra-High Molecular Weight DNA Extraction Protocol V1.4, Circulomics). Blood samples were split based on input requirements for each kit and processed according to the manufacturer’s protocol. Nucleic acid extracts were then quantified by Qubit 4.0 using the Broad Range dsDNA kit (Q32853, ThermoFisher), and for nucleic acid purity using the Nanodrop 2000 (ThermoFisher Scientific). HMW-DNA (High Molecular Weight DNA) was visualized using Pulsed Field Gel Electrophoresis (PFGE) on a Blue Pippen Pulse, set on 70V for 20 hours at room temperature. Samples were stored a −20°C until quantified for sequencing library preparation.

### ONT library preparation and sequencing

DNA from the Nanobind CBB Big DNA kit and the MagMax Core NA kit for both PAXgene and ‘purple top’ EDTA tubes were combined to create a single sample for Oxford Nanopore Technologies library preparation. Half of this sample was used in the Short Read Eliminator Kit (SS-100-101-01, Circulomics, Inc., MD, USA) to test the effect of size-selection on read N50, resulting in a size-selected sample. The size-selected and non-size-selected samples were then split between the Rapid Sequencing Kit (SQK-RAD004, Oxford Nanopore Technologies) and the Ligation Sequencing Kit (SQK-LSK109, Oxford Nanopore Technologies) to test the effect of library preparation on read N50, resulting in a total of four unique libraries. Each library was then loaded onto an R9.4.1 flow cell and sequenced in parallel on the ONT GridION platform. It was determined that size-selection did not have the desired effect of increasing read N50, and four additional non size-selected libraries were prepared (two SQK-RAD004 and two SQK-LSK109) to achieve a target depth of at least 20x. The output of all eight flow cells produced a combined total of approximately 22.7x depth.

### 10X Genomics linked-read sequencing and assembly

For the 10X Genomics assembly, high molecular weight genomic DNA was isolated from whole blood stored in the PAXgene proprietary media using the Nanobind CBB Big DNA kit (Circulomics, Inc., MD, USA) and short fragments filtered out using the Circulomics Short Read Eliminator kit. Genomic DNA concentration and purity were assessed with a Qubit 2.0 Fluorometer (ThermoFisher Scientific, MA, USA) and NanoDrop 2000 spectrophotometer (ThermoFisher Scientific, MA, USA). Capillary electrophoresis was carried out using a Fragment Analyzer (Agilent Technologies, CA, USA) to ensure that the isolated DNA had a minimum molecule length of 40 kb. Genomic DNA was diluted to approximately 1.2 ng/μl and libraries were prepared using Chromium Genome Reagents Kits Version 2 and the 10X Genomics Chromium Controller instrument fitted with a micro-fluidic Genome Chip (10X Genomics, CA, USA). DNA molecules were captured in Gel Bead-In-Emulsions (GEMs) and nick-translated using bead-specific unique molecular identifiers (UMIs; Chromium Genome Reagents Kit Version 2 User Guide) and size and concentration determined using an Agilent 2100 Bioanalyzer DNA 1000 chip (Agilent Technologies, CA, USA). Libraries were then sequenced on an Illumina NovaSeq 6000 System following the manufacturer’s protocols (Illumina, CA, USA) to produce >95x read depth using paired-end 150 bp reads. The reads were assembled into phased pseudo-haplotypes using Supernova Version 2.0 (10X Genomics, CA, USA).

### Genome assembly

As discussed above, two sequencing platforms were employed to sequence and assemble the yellow Labrador Retriever mixed breed *Canis lupus familiaris* phased reference genome; HMW sequencing using R9.4.1 flow cells on ONT’s GridION platform, and 10X Genomics linked-read sequencing on Illumina’s NovaSeq platform. The *de novo* assembly workflow (Figure 6) starts with generating an overlapping read file from all ONT data using minimap2 (version 2.15-r911-dirty) (H. Li 2018). These super-contiguous sequences and the original input read file were then assembled using miniasm (version 0.3-r179) (H. Li 2018). In order to find the best initial assembly for polishing and scaffolding, a range of miniasm parameter combinations were executed as part of this step, and each resulting assembly evaluated for total contig count and length. A five feature parameter space for miniasm was explored, yielding 144 unique parameter tests (see Figure x for specific values used for parameters m[3x], i[3x], s[4x], I[2x], and e[2x]).

**Figure 6.**
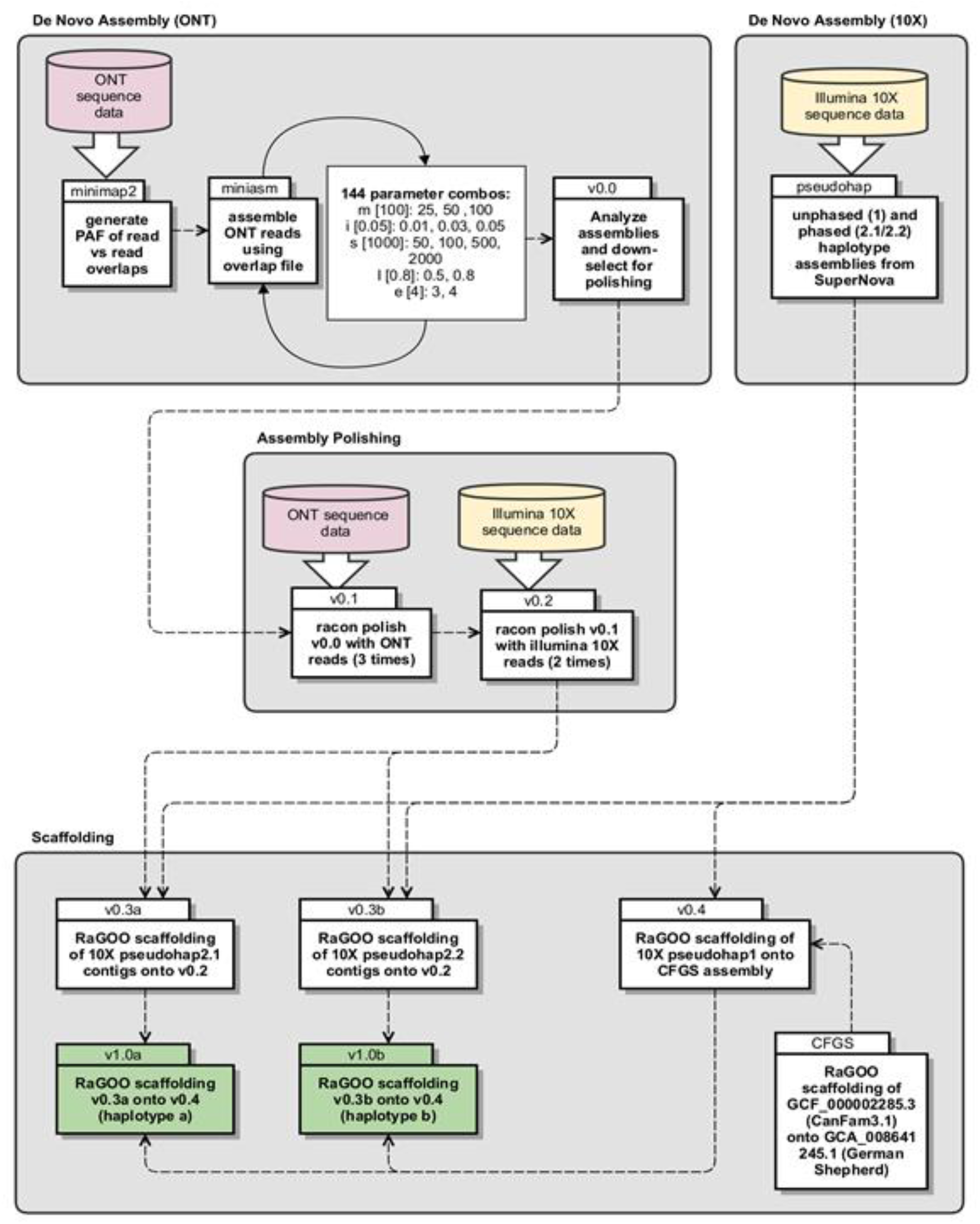
Diagram of phased assembly pipeline. Divided into four primary sections: De Novo Assembly (ONT), De Novo Assembly (10X), Assembly Polishing, and Scaffolding.

After assembly down-selection (v0.0, see Results for specific parameter set), the raw contig correction by rapid assembly methods tool Racon (version v1.4.3) was used for polishing; three rounds with ONT reads (v0.1) followed by two rounds with Illumina 10X reads (v0.2) (Vaser et al. 2017). The read QC tool cutadapt (version 2.5) was used to clip the first 22 bps containing the GEM barcode from the Illumina 10X reads prior to use as polishing input (Martin 2011). Additionally, a base call quality threshold of Phred 20 and a minimum length of 50 bp were used during cutadapt QC processing. In order to produce phased haplotypes, the SuperNova pseudohap2.1 and 2.2 contig sets were scaffolded separately onto v0.2, producing v0.3a and b, respectively (Table 7). The fast and accurate reference-guided scaffolding tool RaGOO (version v1.1) was used to accomplish all scaffolding (Alonge et al. 2019). Alongside polishing and pseudohap phasing of the ONT scaffolds, CanFam3.1 (GCF_000002285.3) was scaffolded onto the newly assembled German Shepherd genome (GCA_008641245.1) (Field et al. 2020) because the latter provides superior chromosomal context for the more fragmented but highly annotated CanFam3.1 genome (CFGS). Next, the unphased SuperNova pseudohap1 contigs were scaffolded onto the CFGS assembly to correct for potential structural variation between breeds, and more accurately reflect the structure of the Labrador Retriever breed (v0.4). Lastly, a final phased v1.0a and b assembly was produced by scaffolding v0.3a and b onto v0.4. Assembly statistics were calculated using QUAST-LG (version v5.0.2), and genome completeness was assessed using BUSCO (version v3, Benchmarking sets of Universal Single-Copy Orthologs) with the mammalia_odb9 dataset (https://busco.ezlab.org/datasets/mammalia_odb9.tar.gz) (Mikheenko et al. 2018; Simão et al. 2015).

**Table 7.** Versions of Yella dog genome assembly. Starting with v0.0, the assembly from miniasm parameter set: m100, i0.05, s500, I0.8, e3 (if not listed, default value was used). The RaGOO generated CFGS assembly is the primary reference used for chromosomal scale

| Version | Description |
| --- | --- |
| CFGS | RaGOO of CanFam3.1 onto German Shepherd scaffolds |
| 0.0 | raw assembly from miniasm |
| 0.1 | v0.0 + 3x racon polishing with ONT reads |
| 0.2 | v0.1 + 2x racon polishing with illumina 10X reads |
| 0.3a | RaGOO of 10X pseudohap2.1 contigs onto v0.2 |
| 0.3b | RaGOO of 10X pseudohap2.2 contigs onto v0.2 |
| 0.4 | RaGOO of JHMI 10X psuedohap scaffolds onto CFGS scaffolds |
| 1.0a | RaGOO of v0.3a onto v0.4 |
| 1.0b | RaGOO of v0.3b onto v0.4 |

### Alignment and variant calling

Reads from SRA were aligned to the three canine reference genomes shown in Figure 5 using default parameter settings for the graph-based aligner HISAT2 (Kim et al. 2019). Secondary and supplementary alignments were then filtered using samtools with parameters “-F0×4 -F0×100 -F0×800” (Li et al. 2009). Variant calling was performed using default parameters for “bcftools mpileup” and “bcftools call”, then filtering out variant calls with QUAL less than 30 (Li 2011).

## DATA ACCESS

The sequence read data and assemblies generated in this study have been submitted to the NCBI BioProject database (https://www.ncbi.nlm.nih.gov/bioproject/) under accession number PRJNA610592. All samples used in this study are under BioSample SAMN14279123. The primary haplotype FASTAs are under BioProject PRJNA610232 and differentiated from the alternative haplotype with an ‘a’ at the end of header names excluding the MT header (40 sequences, MT included, GenBank accessions CP050567.1 - CP050606.1). The alternative haplotype FASTAs are under BioProject PRJNA610230 and differentiated from the primary with a ‘b’ at the end of header names (39 sequences, MT ommited, GenBank accessions CP050607.1 - CP050645.1).

## ACKNOWLEDGEMENTS

Funding for this project was provided by the Department of Homeland Security (DHS) Science and Technology Directorate (S&T), Contract No. 70RSAT19CB0000002. Karen Meidenbauer, DVM, is acknowledged for her technical leadership and expertise, as well as her participation drawing blood, which was transported via shippable pelican case with all lab equipment, reagents, and samples. David Deglau and Michael House are gratefully acknowledged for project and program management, respectively. Jody Proescher is acknowledged for her critical review of and editorial feedback for the manuscript.

## AUTHOR CONTRIBUTIONS

RP performed all bioinformatics analysis and generated all figures and tables, and large portions of the manuscript. EF and KV performed wet lab studies including high molecular weight DNA extractions, library preparation, and nanopore sequencing. DM performed all 10X Genomics and Illumina NovaSeq experiments and bioinformatics analysis. AS funded the 10X Genomic and NovaSeq experiments, and contributed intellectually to the study integration of Illumina and nanopore data. CB proposed and established the initial study, provided scientific leadership, and contributed large portions of the manuscript.

## Acronyms

bp: base pair
BUSCO: Benchmarking sets of Universal Single-Copy Orthologs
CFGS: scaffold of CanFam3.1 (GCF_000002285.3) on German Shepherd genome (GCA_008641245.1)
EDTA: ethylenediaminetetraacetic acid
Gb: gigabase
GWAS: Genome Wide Association Study
HMW-DNA: High Molecular Weight DNA
kb: kilobase
NHGRI: National Human Genome Research Institute
ONT: Oxford Nanopore Technologies
PCE: Phenol-Chloroform Extraction
PFGE: Pulsed Field Gel Electrophoresis
SNP: Single Nucleotide Polymorphism
WGS: Whole Genome Sequencing

## SUPPLEMENTAL MATERIAL

**Figure S1.**
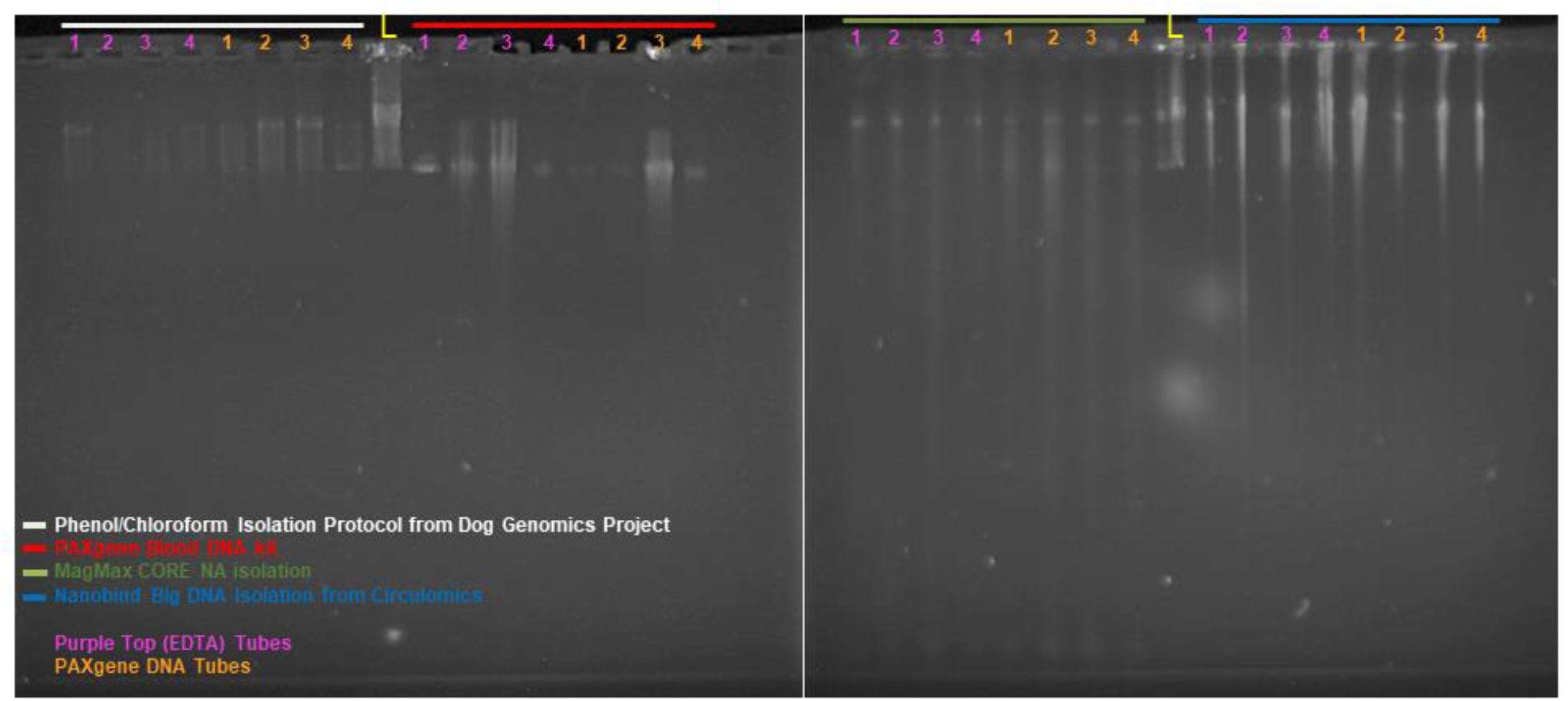
PFGE visualization of HMW-DNA extracted from four different extraction methods (PCE. PAXgene MagMax, and Nanobind) using blood stored in two different preservation agents (PAXgene and EDTA). Samples are from four dogs (numbered across the top 1-4). The lambda ladder is in the middle lane of each gel. indicated by a yellow V (48.5Kb - 1Mb. 18 bands at 48.5Kb steps). The PAXgene extraction kit is the only kit that failed to yield HMW-DNA. PFGE run at 70V for 20 hours.

**Figure S2.**
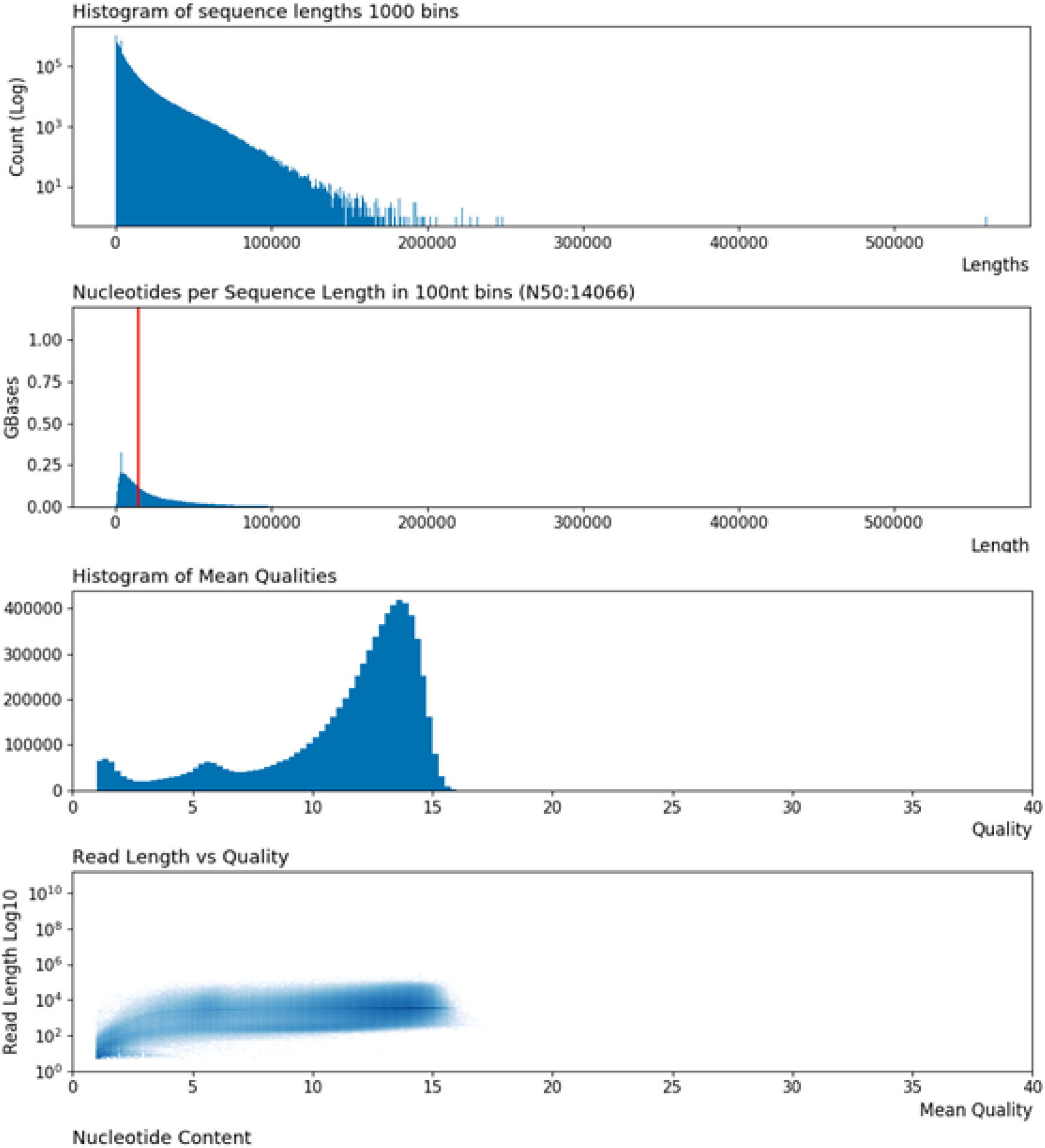
Sequence length and base call quality distributions of combined read data from all eight ONT flow cells used to generate at least 2Ox depth across the approximately 2.3Gb canine genome.

**Figure S3.**
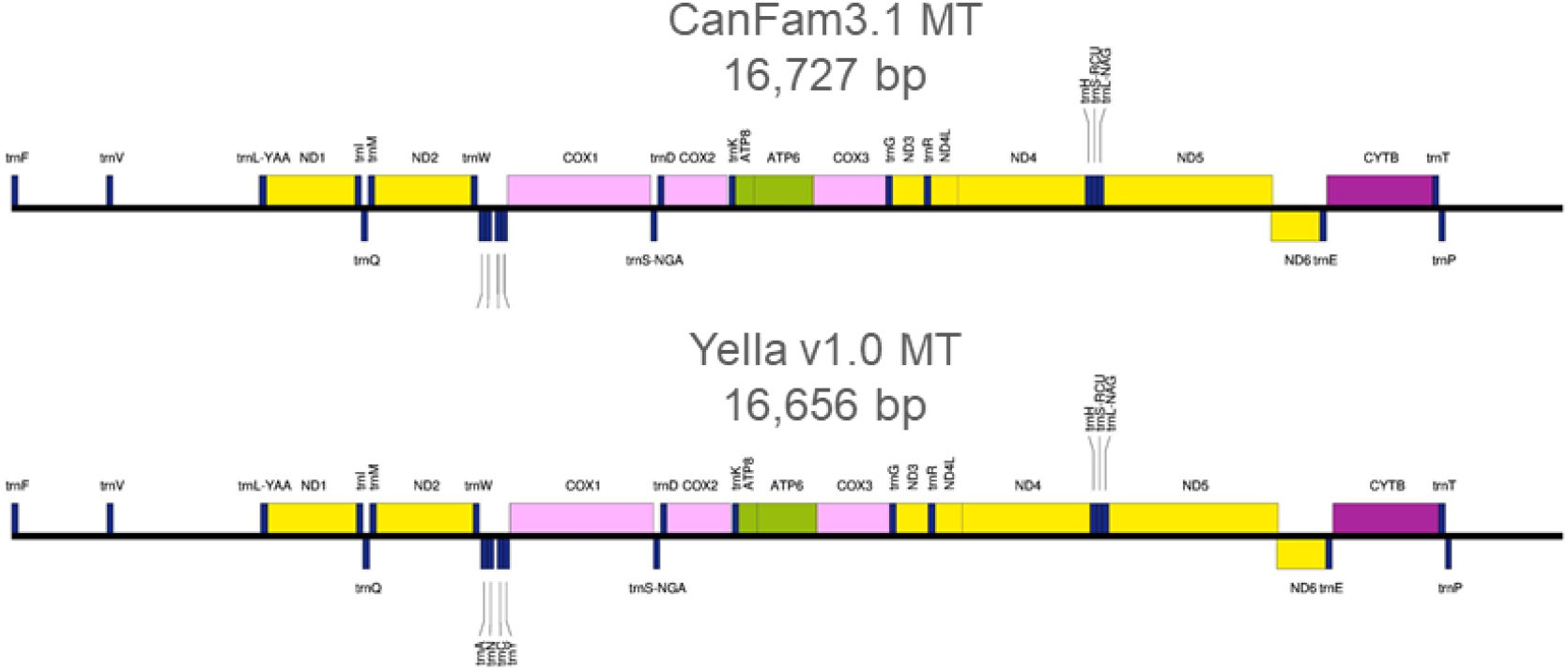
Sequence annotation maps of refseq canine mitochondrial sequence (top) and Yelia v1.0 mitochondrial sequence (bottom). Maps generated from GeSeq’s Chlorobox annotation and visualization web tool (reference below). Alignment of Yelia MT to refseq MT reveals 3 bps of insertions and 74 bps of deletion (alignment CIGAR: 2678M1I7233M2I6327M50D36M24D379M). Needleman-Wunsch pairwise alignment results 99.41 identity and similarity between these MT sequences.

**Table S1.** Supplementary data from two storage and four nucleic acid (NA) extraction kits. Blood was preserved from four dogs (including Yella, Dog ID #7) using two different storage agents, then NA isolated using four different extraction kits. Subsets of this data were used in Tables 2 and 3. DNA 260/280 ratio, ~1.8 is considered ‘pure’ for DNA, ~2.0 is considered ‘pure’ for RNA. Expected 260/230 values are commonly in the range of 2.0–2.2.

| Sample ID | Dog ID | Storage Agent | Isolation Kit | Isolation Volume (uL) | Extracted NA Concentration (ng/uL) | Total Extracted NA (ng) | Total NA Normalized to Kit Input Volume (ng) | NA Quality (260/280) | NA Quality (260/230) | HMW DNA Yielded? |
| --- | --- | --- | --- | --- | --- | --- | --- | --- | --- | --- |
| 12928 | 5 | purple top tube (EDTA) | PCE | 1000 | 3.7 | 3700 | 3.7 | 2.4 | 4.6 | yes |
| 12929 | 5 | purple top tube (EDTA) | PAXgene | 1000 | 0.36 | 360 | 0.36 | 0.55 | 0.08 | no |
| 12930 | 5 | purple top tube (EDTA) | Nanobind | 100 | 19.5 | 1950 | 19.5 | 1.89 | 1.25 | yes |
| 12931 | 5 | purple top tube (EDTA) | Magmax | 90 | n/a | - | - | - | - | - |
| 12932 | 5 | PAXgene (proprietary) | PCE | 1000 | 1.11 | 1110 | 1.11 | 6.11 | -3.11 | yes |
| 12933 | 5 | PAXgene (proprietary) | PAXgene | 1000 | 0.18 | 180 | 0.18 | 0.47 | 0.09 | no |
| 12934 | 5 | PAXgene (proprietary) | Nanobind | 100 | 8.32 | 832 | 8.32 | 1.93 | 0.89 | yes |
| 12935 | 5 | PAXgene (proprietary) | Magmax | 90 | 1.52 | 136.8 | 1.52 | 1.74 | 0.2 | yes |
| 12938 | 6 | purple top tube (EDTA) | PCE | 1000 | 0.28 | 280 | 0.28 | -2.3 | -0.43 | yes |
| 12939 | 6 | purple top tube (EDTA) | PAXgene | 1000 | 4.44 | 4440 | 4.44 | 2.21 | 9.32 | no |
| 12940 | 6 | purple top tube (EDTA) | Nanobind | 100 | 54 | 5400 | 54 | 1.84 | 1.16 | yes |
| 12941 | 6 | purple top tube (EDTA) | Magmax | 90 | 22.2 | 1998 | 22.2 | 1.56 | 0.23 | yes |
| 12942 | 6 | PAXgene (proprietary) | PCE | 1000 | 5.46 | 5460 | 5.46 | 2.3 | 3.8 | yes |
| 12943 | 6 | PAXgene (proprietary) | PAXgene | 1000 | 0.2 | 200 | 0.2 | -0.69 | -0.01 | no |
| 12944 | 6 | PAXgene (proprietary) | Nanobind | 100 | 18.7 | 1870 | 18.7 | 1.88 | 1.85 | yes |
| 12945 | 6 | PAXgene (proprietary) | Magmax | 90 | 11.48 | 1033.2 | 11.48 | 1.64 | 0.27 | yes |
| 12948 | 7 | purple top tube (EDTA) | PCE | 1000 | 0.38 | 380 | 0.38 | 5.21 | -1.68 | yes |
| 12949 | 7 | purple top tube (EDTA) | PAXgene | 1000 | 10.8 | 10800 | 10.8 | 1.98 | 6.79 | no |
| 12950 | 7 | purple top tube (EDTA) | Nanobind | 100 | 35.3 | 3530 | 35.3 | 1.84 | 1.69 | yes |
| 12951 | 7 | purple top tube (EDTA) | Magmax | 90 | 2.63 | 236.7 | 2.63 | 1.62 | 0.26 | yes |
| 12952 | 7 | PAXgene (proprietary) | PCE | 1000 | 6.37 | 6370 | 6.37 | 2.2 | 3.79 | yes |
| 12953 | 7 | PAXgene (proprietary) | PAXgene | 1000 | 6.4 | 6400 | 6.4 | 2.38 | -5.4 | no |
| 12954 | 7 | PAXgene (proprietary) | Nanobind | 100 | 11.1 | 1110 | 11.1 | 1.87 | 2.05 | yes |
| 12955 | 7 | PAXgene (proprietary) | Magmax | 90 | 2.03 | 182.7 | 2.03 | 1.66 | 0.35 | yes |
| 12958 | 8 | purple top tube (EDTA) | PCE | 1000 | 0.75 | 750 | 0.75 | 1.7 | 3.37 | yes |
| 12959 | 8 | purple top tube (EDTA) | PAXgene | 1000 | 0.7 | 700 | 0.7 | 2.17 | -0.19 | no |
| 12960 | 8 | purple top tube (EDTA) | Nanobind | 100 | 8.13 | 813 | 8.13 | 1.84 | 0.92 | yes |
| 12961 | 8 | purple top tube (EDTA) | Magmax | 90 | 2.33 | 209.7 | 2.33 | 1.53 | 0.33 | yes |
| 12962 | 8 | PAXgene (proprietary) | PCE | 1000 | 1.2 | 1200 | 1.2 | 2.04 | 2.59 | yes |
| 12963 | 8 | PAXgene (proprietary) | PAXgene | 1000 | 0.59 | 590 | 0.59 | -2.48 | -0.1 | no |
| 12964 | 8 | PAXgene (proprietary) | Nanobind | 100 | 34.4 | 3440 | 34.4 | 1.95 | 1.47 | yes |
| 12965 | 8 | PAXgene (proprietary) | Magmax | 90 | 1.53 | 137.7 | 1.53 | 1.61 | 0.3 | yes |

**Table S2.**
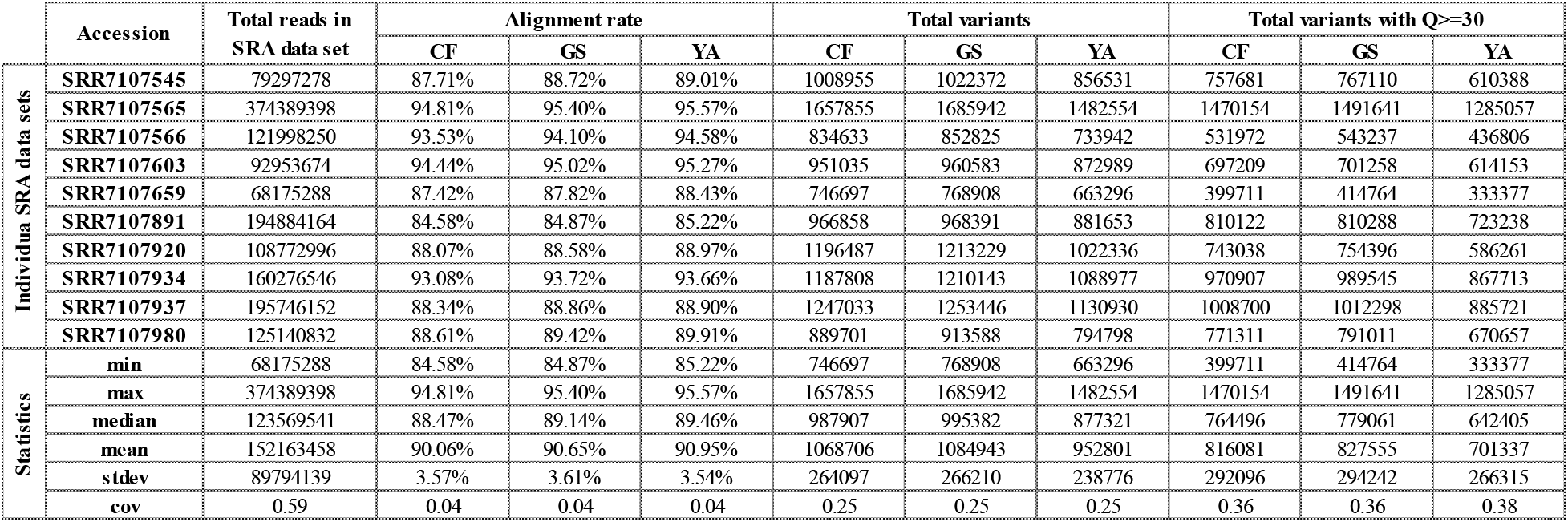
Alignment rates and total variants of ten Labrador Retriever Illumina sequence read data sets from SRA, with additional metrics and summary statistics. CF, Boxer (CanFam3.1), GCF_000002285.3; GS, German Shepherd, GCA_008641245.1; YA, Labrador Retriever (Yella_v1.0), CP050567.1 - CP050606.1

